# Lipopolysaccharide induces placental mitochondrial dysfunction by reducing MNRR1 levels via a TLR4-independent pathway

**DOI:** 10.1101/2021.11.06.467519

**Authors:** Neeraja Purandare, Yusef Kunji, Yue Xi, Roberto Romero, Nardhy Gomez-Lopez, Andrew Fribley, Lawrence I. Grossman, Siddhesh Aras

## Abstract

Mitochondria play a key role in the growth and development of the placenta, an organ essential for pregnancy in eutherian mammals. Mitochondrial dysfunction has been associated with pregnancy pathologies. However, the mechanisms whereby placental mitochondria sense inflammatory signals at a cellular and mechanistic level are unknown. Mitochondrial Nuclear Retrograde Regulator 1 (MNRR1) is a bi-organellar protein responsible for optimal mitochondrial function to achieve energy and redox homeostasis. In addition, MNRR1 also is required for optimal induction of cellular stress-responsive signaling pathways such as the mitochondrial unfolded protein response (UPR^mt^). Here, in a lipopolysaccharide-induced model of placental inflammation, we show that MNRR1 levels are reduced in placental tissues and cell lines. Reduction in MNRR1 is associated with mitochondrial dysfunction and enhanced oxidative stress along with activation of pro-inflammatory signaling. Mechanistically, we uncover a non-conventional pathway independent of Toll-like receptor 4 (TLR4) that results in a specific ATM kinase-dependent threonine phosphorylation and activation of a mitochondrial protease, YME1L1, degrading MNRR1. Furthermore, enhancing MNRR1 levels in placental cells either genetically or with specific activators abrogates the bioenergetic defect and induces an anti-inflammatory phenotype, suggesting that MNRR1 is upstream of the mitochondrial dysfunction observed in our model. Reduction in MNRR1 levels is a generalized phenomenon observed in cells under an inflammatory stimulus. We therefore propose MNRR1 as a novel anti-inflammatory therapeutic target in pathologies associated with placental inflammation.

## Introduction

Spontaneous preterm birth – the birth of a baby before 37 weeks of gestation – is the leading cause of neonatal mortality and morbidity worldwide [1, 2]. Spontaneous preterm birth is preceded by preterm labor, a syndrome of multiple etiologies including local and systemic inflammation [3], which results from the immune activation triggered by microbes invading the amniotic cavity [4] or danger signals derived from cellular necrosis or stress [5]. Most research has focused on investigating the inflammatory pathways taking place in the intra-amniotic cavity containing the placenta; however, the role of mitochondria acting as a sensor of cellular stress has been less investigated.

Under healthy conditions, mitochondria are required to generate energy for cellular functioning in the form of ATP. This process is fine-tuned to respond to stress signals by slowing ATP production and activating immune response pathways such as by generating reactive oxygen species (ROS) [6] and other biological processes [6–8]. The role of placental mitochondria has only recently been reported in normal gestation [9, 10]. Yet, mitochondrial dysfunction has also been associated with pregnancy complications including preeclampsia [11, 12], intrauterine growth restriction [13], maternal adiposity [14, 15], gestational diabetes [16], and spontaneous preterm birth [17], all of which are associated with inflammatory responses [18–25]. Specifically, these studies have shown that some electron transport chain (ETC) subunits, chaperones, ROS-scavenging enzymes, and mitochondrial DNA (mtDNA) levels are altered in such complicated pregnancies. However, the mechanisms whereby placental mitochondria sense inflammatory signals at a cellular and mechanistic level are not clearly known.

We previously showed that MNRR1 (CHCHD2; AAG10) regulates mitochondrial function by acting in two compartments – the mitochondria and the nucleus [26–29]. Mitochondrial MNRR1 interacts with complex IV (cytochrome *c* oxidase; COX) of the ETC to regulate oxygen consumption and can alter apoptosis by interacting with Bcl-xL [30]. In the nucleus, MNRR1 regulates the transcription of numerous genes including subunits of ETC complexes, ROS scavenger genes, and proteins that regulate mitochondrial proliferation [27, 31, 32]. Here, we have characterized the role of MNRR1 *in vivo* using placental tissues from a murine model of lipopolysaccharide (LPS)-induced preterm birth and *in vitro* using cultured trophoblast (i.e., placental) cell lines. We find that LPS reduces MNRR1 levels in placental tissue as well as in trophoblast cell lines. We then went on to identify a novel pathway that results in MNRR1-dependent mitochondrial dysfunction, thereby uncovering potential therapeutic targets. Taken together, our work shows that MNRR1 plays a protective role by not only activating mitochondria but also by inducing an anti-inflammatory response to ameliorate the deleterious effects of placental inflammation.

## Results

### Decreased MNRR1 impairs mitochondrial function in human placental cells *in vitro*

MNRR1 modulates several mitochondrial functions including oxygen consumption, ATP production, and generation of ROS [27]. To analyze these effects in a human system *in vitro*, we generated a placental cell culture model of LPS-induced inflammation using the trophoblast cell line HTR8/SVNeo (HTR). Since inflammation suppresses mitochondrial function [33–38], we first determined the effect of inflammation on MNRR1 in these placental cells. When we measured the basal oxygen consumption rate (OCR) in intact trophoblast cells treated with LPS, we found an ~30% decrease (**Fig. 1A**). We wondered whether this decreased OCR affects ATP levels and found an ~18% decrease in total cellular ATP (**Fig. 1B**). Furthermore, both total intracellular ROS and mitochondrial ROS were increased (**Fig. 1C**). Similar findings were previously made in immune cells [6] and for total ROS in another trophoblast cell line, Sw.71 [39]. This decrease in OCR can be completely rescued (and enhanced) by overexpressing WT-MNRR1 (**Fig. 1D**). The decrease in OCR is consistent with inhibition also at the protein level (**Fig. 1E, Supplementary Figure 1A)**. To further link MNRR1 levels with mitochondrial function, we inhibited MNRR1 expression pharmacologically (Clotrimazole; **Supplementary Figure 2A**), which sensitized cells to the effect of LPS on OCR (**Fig. 1F (left), bars 3 and 4**) and an activator (Nitazoxanide; **Supplementary Figure 2A**), which prevented these effects (**Fig. 1F (left), bars 5 and 6**). We also examined the effects of MNRR1 inhibition and activation on an inflammatory marker by assessing JNK phosphorylation (**Fig. 1E**). Chemical inhibition of MNRR1 acts similarly to LPS mediated inflammation in increasing JNK phosphorylation (**Fig. 1F (right)**). Importantly, chemical activation of MNRR1 by Nitazoxanide prevents this increase even after LPS-treatment (**Fig. 1F (right)**), suggesting that such activators could be repurposed therapeutically to treat placental inflammation.

**Figure 1:**
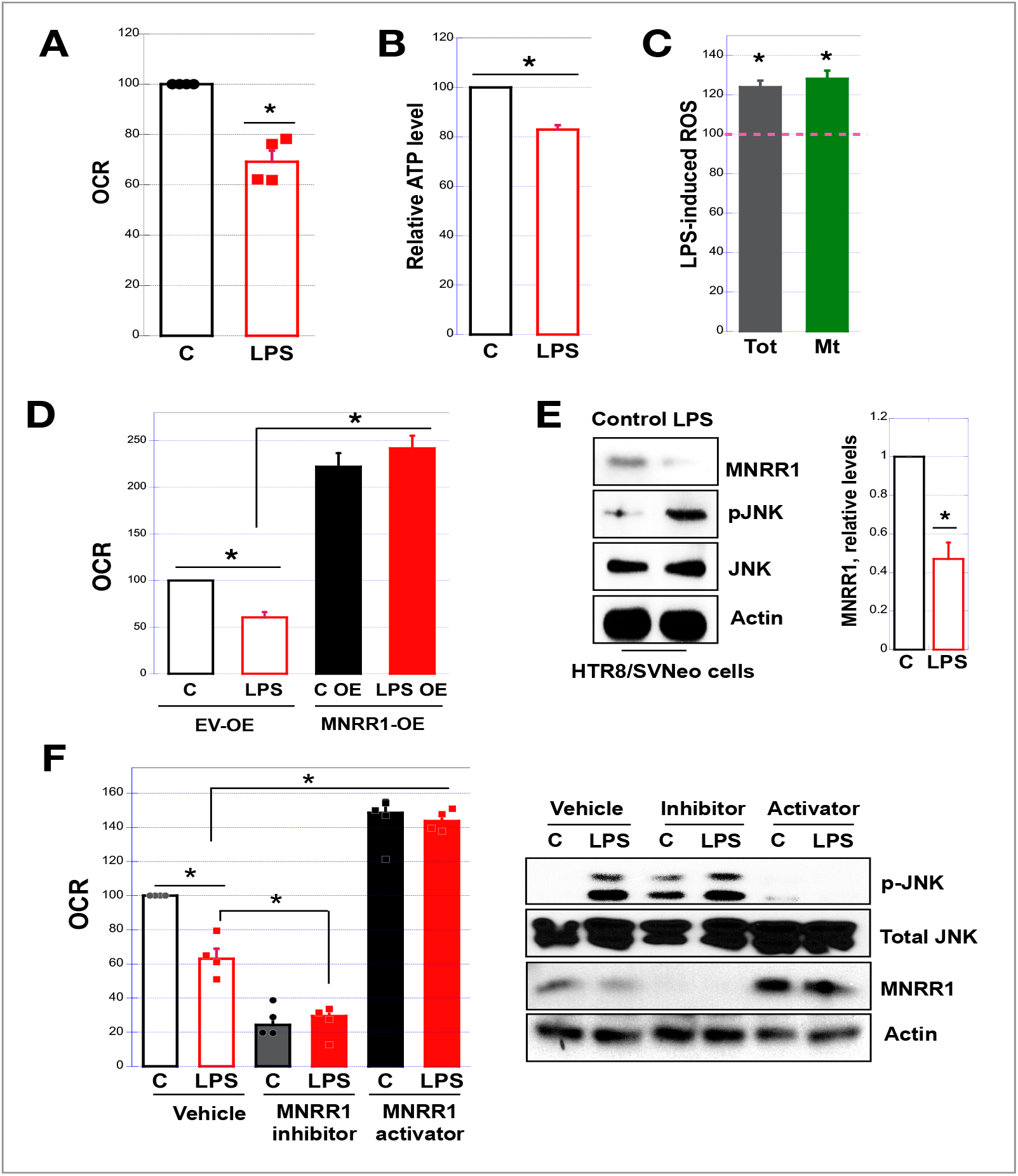
LPS decreases MNRR1 levels and impairs mitochondrial function in human placental cells, defects that can be rescued by increasing MNRR1 expression. All LPS treatments in cultures cells are at 500 ng/mL for 24 h unless indicated otherwise. In all figures: *, p <0.05; **, p < 0.005. **(A)** Intact cellular oxygen consumption in the HTR cells treated with control (water) or LPS. Data are represented as oxygen consumption relative to control set to 100%. **(B)** Equal numbers of HTR cells were plated in a 96-well plate and ATP levels were measured as in Experimental Procedures. **(C)** HTR cells were treated with control (water) or LPS and ROS levels were measured using CM-H2DCFDA (total, Ex: 485 nm/Em: 527 nm) or MitoSOX Red (mitochondrial, Ex: 510 nm/ Em: 580 nm). **(D)** Intact cellular oxygen consumption in HTR cells overexpressing EV or MNRR1 and treated with control or LPS. Data are represented as oxygen consumption relative to EV-control set to 100%. **(E)** *Left*, Equal amounts of HTR cells treated with control or LPS were separated on an SDS-PAGE gel and probed for MNRR1, phospho-JNK, and total JNK levels. Actin was probed as loading control. *Right*, The graph represents MNRR1 levels relative to Actin. **(F)** *Left*, Intact cellular oxygen consumption in HTR cells treated with Vehicle (DMSO), MNRR1 inhibitor (Clotrimazole (C), 10 μM), or MNRR1 activator (Nitazoxanide (N), 10 μM) with control (water) or LPS (500 ng/mL) for 24 h. Data are expressed relative to EV-control set to 100%. *Right*, Pooled lysates from OCR measurement were separated on an SDS-PAGE gel and probed for MNRR1, phospho-JNK, and total JNK levels. Actin was probed as a loading control.

### MNRR1 levels are reduced in a murine bacterial endotoxin model of placental inflammation *in vivo*

To assess the *in vivo* relevance of our observations in cultured placental cells, we utilized a mouse model of LPS-induced inflammation. LPS treatment is known to induce a high rate (~80%) of preterm labor and birth [40]. Our examination of MNRR1 protein levels in placental lysates from LPS-treated mice showed them to be significantly decreased (**Fig. 2A**). We then immunostained the placental tissue from LPS injected mice and found MNRR1 protein levels to be reduced compared to those injected with phosphate buffered saline (controls) (**Fig. 2B**). Given the reduced MNRR1 levels after LPS treatment, we analyzed whether transcript levels of MNRR1 were also reduced but found them to be unchanged (**Supplementary Fig. 1B**), suggesting a post-translational effect.

**Figure 2:**
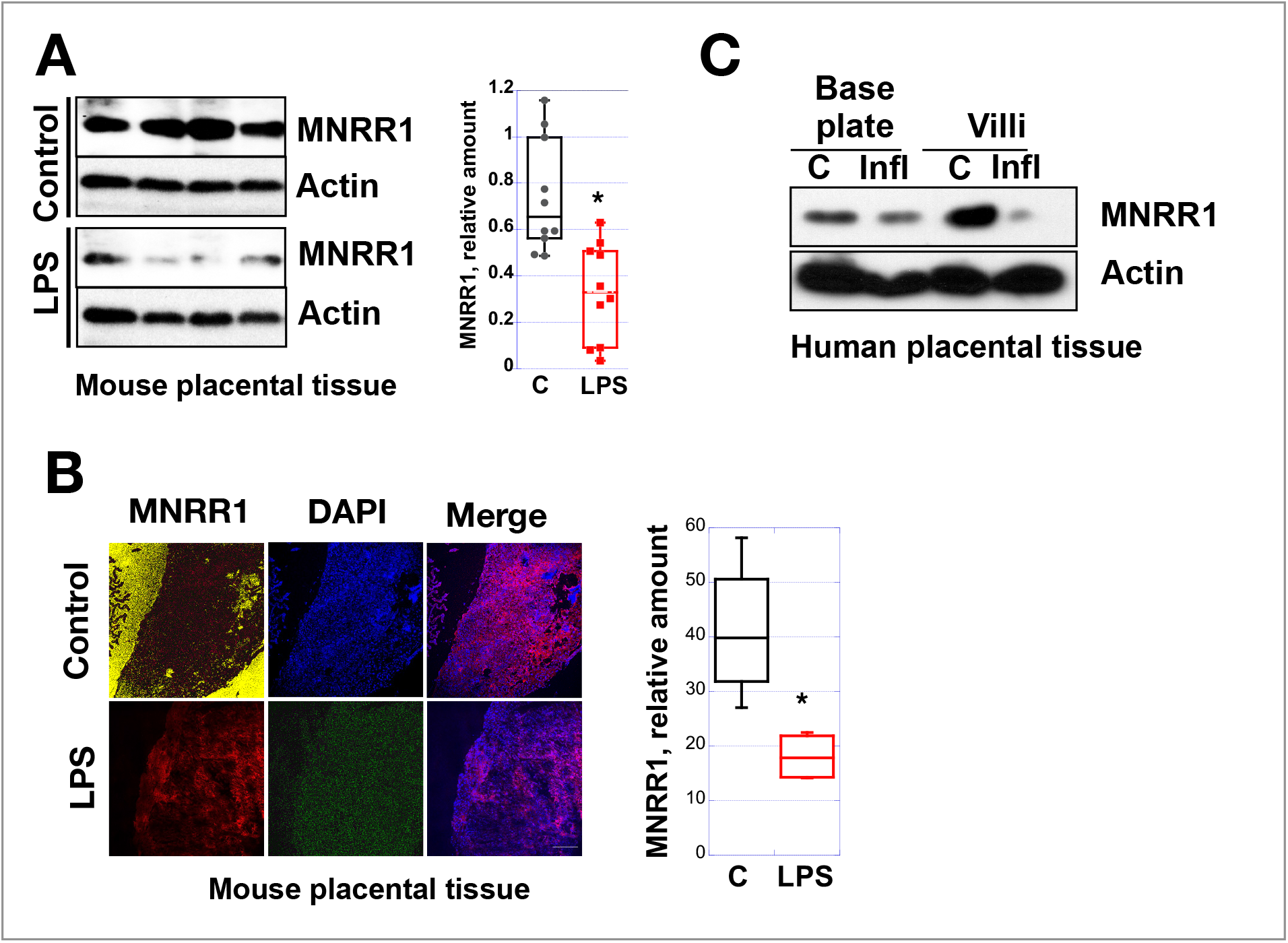
MNRR1 protein levels are reduced *in vivo* in placental inflammation. **(A)** *Left*, Placental lysates from control (PBS) versus LPS (intraperitonially) injected mice were separated on an SDS-PAGE gel and probed for MNRR1 levels. Actin was used as a loading control. *Right*, MNRR1 levels relative to Actin with one point for each animal. **(B)** *Left*, Placental tissue sections from control (PBS) versus LPS injected mice analyzed using immunofluorescence staining. *Right*, Relative MNRR1 fluorescence is shown. **(C)** Equal amounts of human placental lysates from an individual without inflammation and one with systemic inflammation were separated on SDS-PAGE gel and probed for MNRR1. Actin was probed as a loading control.

Since the intraperitoneal injection of LPS in pregnant mice is considered to model a systemic maternal inflammatory response [40, 41], we tested a human placental tissue sample from women with clinical chorioamnionitis [42] who delivered preterm. In this sample, we found MNRR1 levels considerably reduced in the villous layer of the placenta and moderately so in the base plate (**Fig. 2C**). Mirroring the *in vivo* data from mice, MNRR1 was reduced at the protein level in HTR cells (**Fig. 1E**) but not at transcript levels (**Supplementary Fig. 1C and 1D**). We also measured JNK phosphorylation [43, 44], and found increased activation (**Fig. 1E**). Our data with both mouse and human placental samples thus suggest that MNRR1 reduction occurs in response to maternal systemic inflammation. We considered whether other cell types reduce MNRR1 levels in response to LPS and examined both placental and non-placental cell lines. We found reduced MNRR1 levels after LPS-treatment in all the cell lines tested, suggesting that MNRR1 reduction is a ubiquitous phenomenon (**Supplementary Fig. 1A**).

### Increased YME1L1 protease reduces mitochondrial MNRR1 levels in human placental cells *in vitro*

Since MNRR1 is a bi-organellar protein that is localized both to the mitochondria and the nucleus [27–29], we determined the effect of LPS on MNRR1 levels in each of the compartments. We found that most of the decrease at the protein level was accounted for by mitochondrial MNRR1 (**Figs. 3A and 3B**), strikingly so when visualized by confocal microscopy (**Fig. 3B**). To investigate how the mitochondrial reduction in MNRR1 takes place, we assessed the protein levels of YME1L1, a mitochondrial intermembrane space (IMS) protease previously shown to be responsible for the turnover of mitochondrial MNRR1 [32]. We found that levels of YME1L1 are increased by LPS treatment (**Fig. 3C**). We then tested whether MNRR1 is reduced in the absence of YME1L1 and found that it is not by using LPS-treated 293 cells from which YME1L1 had been knocked out (YME1L1-KO) (**Supplementary Fig. 3A**). The levels of OMA1, another protease that has been identified to turn over MNRR1 under cellular stress [45], are not increased with LPS treatment (**Supplementary Fig. 3A and B**), thereby suggesting that the upstream inflammatory signaling pathway involves only YME1L1. To further define the role of YME1L1 in regulating MNRR1 levels in mitochondria, we utilized a version of YME1L1 mutated to eliminate protease activity (protease-dead; PD) [46]. In cells overexpressing PD-YME1L1, levels of MNRR1 were again not reduced after LPS treatment (**Fig. 3D**). Moreover, examination of a known substrate of YME1L1 proteolysis, STARD7 [46, 47], showed LPS-stimulated reduction with active YME1L1 but not with PD-YME1L1, like MNRR1 (**Fig. 3D**). Thus, we find that protease YME1L1 levels are increased in cells treated with LPS, thereby reducing the level of MNRR1.

**Figure 3:**
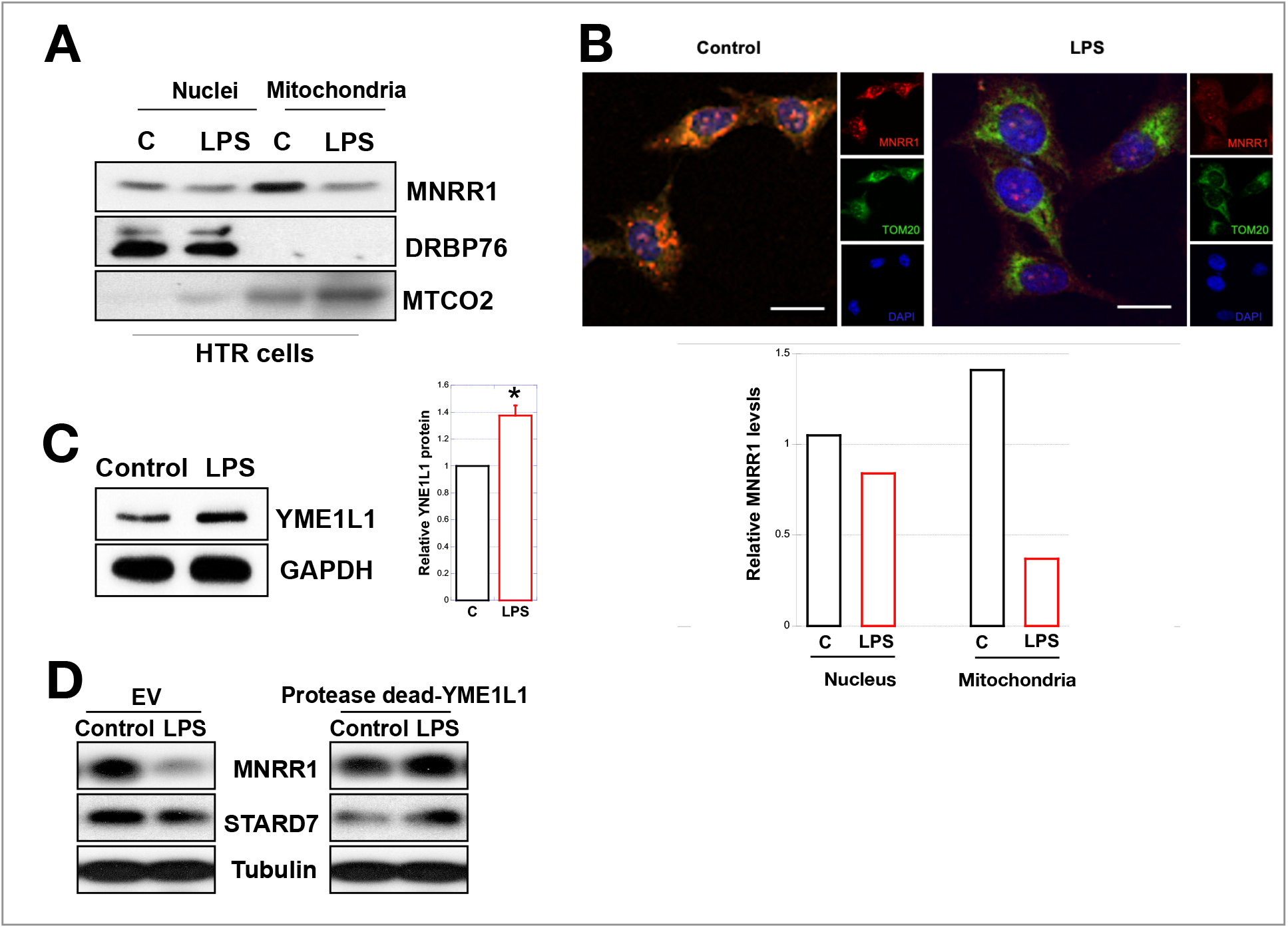
Compartment-specific reduction in MNRR1 levels in human placental cells treated with LPS is mediated by YME1L1 protease. **(A)** Equal amounts HTR cell nuclear and mitochondrial fractions were separated on an SDS-PAGE gel and probed for MNRR1. DRBP76 and MTCO2 were probed to assess purity of fractions. **(B)** HTR cells were treated with control or LPS and immunostained for MNRR1 (red fluorescence). DAPI (blue fluorescence) was stained as a nuclear marker and TOM20 (green fluorescence) as a mitochondrial marker. *Below*, The graph represents MNRR1 levels relative to a compartment-specific control for each treatment. **(C)** *Left*, Equal amounts of HTR cells treated with control or LPS were separated on an SDS-PAGE gel and probed for YME1L1. Actin was probed as loading control. *Right*, The graph represents YME1L1 levels relative to GAPDH. **(D)** HTR cells overexpressing either an empty vector (EV) or a protease-dead (PD) mutant of YME1L1. Cells were treated with control or LPS. Equal amounts of cell lysates were separated on an SDS-PAGE gel and probed for MNRR1 and STARD7. Tubulin was probed as loading control.

### ATM kinase mediated phosphorylation of YME1L1 enhances its stability

The finding that YME1L1 protein levels were increased in LPS-treated placental cells (**Fig. 3C**) whereas transcript levels were unaffected (**Supplementary Fig. 3C**) suggested increased protein stability. We confirmed this finding by carrying out a cycloheximide chase experiment, which showed increased stability after blocking new protein synthesis (**Fig. 4A**). To uncover the basis of the increased stability, we hypothesized a protein modification and thus examined the post-translational profile of YME1L1. Upon treatment with LPS, YME1L1 protein in HTR cells displayed enhanced threonine phosphorylation but not serine or tyrosine phosphorylation (**Fig. 4B, Supplementary Fig. 3D**).

**Figure 4:**
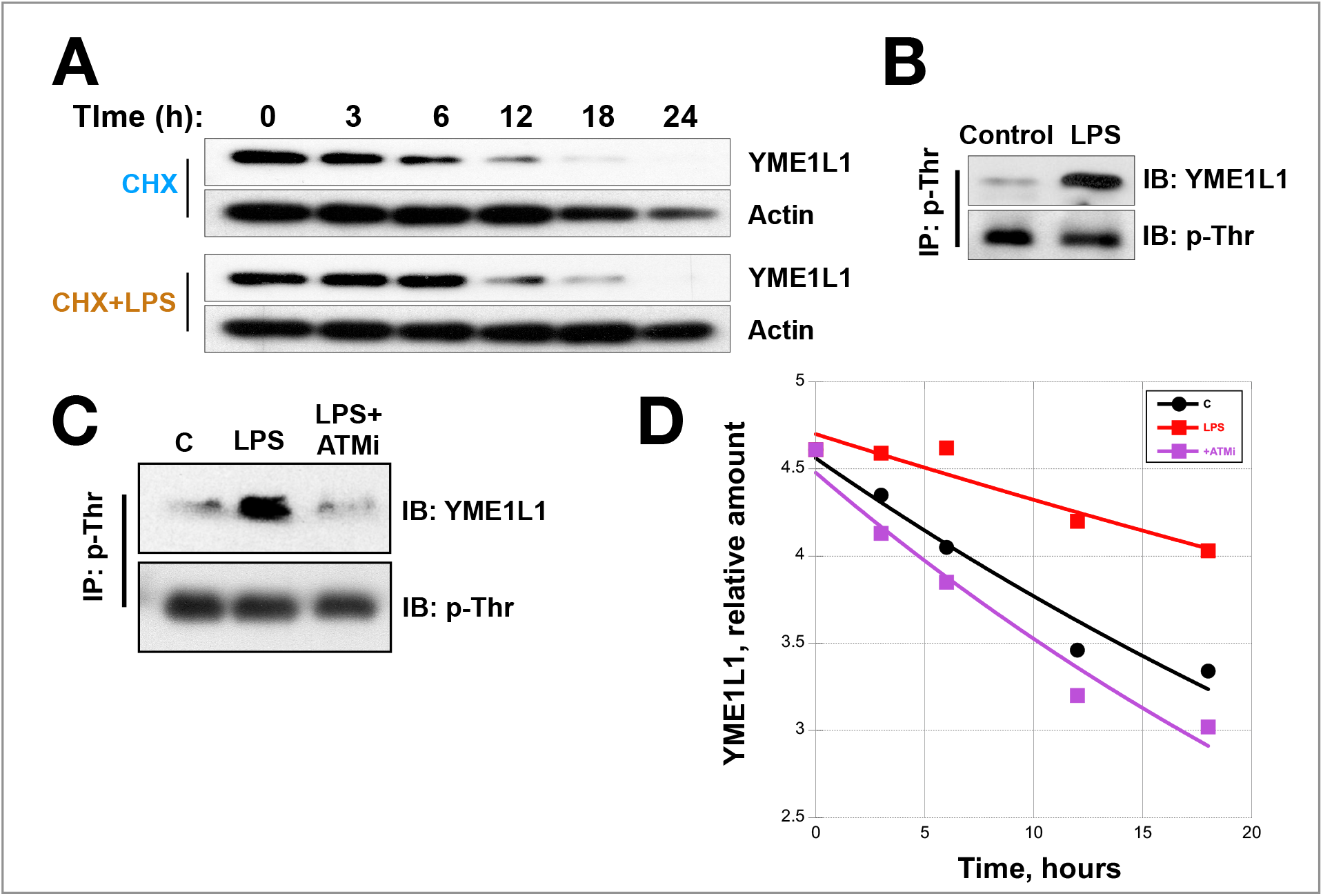
LPS treatment of placental cells increases the stability of YME1L1 via threonine phosphorylation by ATM kinase. **(A)** Equal numbers of HTR cells were plated on a 6-well plate and treated with control or LPS and 100 μg/mL cycloheximide for the durations shown. Equal amounts of cell lysates were separated on an SDS-PAGE gel and probed for YME1L1. Actin was probed as loading control. **(B)** HTR cells were treated with vehicle or LPS and equal amounts of whole cell lysates were used for immunoprecipitation using a phospho-threonine antibody. Equal amounts IP eluates were then probed for YME1L1. Antibody heavy chain (p-Thr) was probed to assess loading. **(C)** HTR cells were treated with control, LPS, or LPS+ATM inhibitor (1 μM) for 24 h. Equal amounts of whole cell lysates were used for immunoprecipitation by phospho-threonine antibody. Equal amounts IP eluates were probed for YME1L1 antibody and heavy chain (p-Thr) was probed to assess loading. **(D)** Graph for YME1L1 levels from HTR cells were treated with control, LPS, or LPS+ATM inhibitor (1 μM) and 100 μg/mL cycloheximide for the durations shown in (A). The amount relative to time = 0 was graphed.

To identify the threonine kinase for which YME1L1 is a substrate, we used Scansite (https://scansite4.mit.edu/4.0/#home), which identified ATM and NEK6 as candidate kinases for YME1L1 under high stringency conditions (**Supplementary Fig. 4A**). Of these, ATM kinase was found to interact with YME1L1 in LPS-treated placental cells (**Supplementary Fig. 4B**) whereas NEK6 kinase did not (**Supplementary Fig. 4C**). To further assess this bioinformatic prediction, we utilized an inhibitor of ATM kinase activity and found that LPS-stimulated threonine phosphorylation of YME1L1 was blocked (**Fig. 4C**). Furthermore, turnover of MNRR1 and YME1L1 substrate STARD7 was also blocked by the same ATM inhibitor (**Supplementary Fig. 4D**). We next asked whether LPS induced threonine phosphorylation of YME1L1 can affected stability of the protease. We found that YME1L1 half-life (8.1 h) is more than doubled by LPS treatment (22.0 h) and that this stabilization is lost when ATM kinase is inhibited (**Fig. 4D**). These results suggest that YME1L1 stability is enhanced upon threonine phosphorylation by ATM kinase in LPS treated placental cells, resulting in increased MNRR1 turnover with subsequent reduction of mitochondrial OCR (**Figs. 1A, 1D, and 1F**), increased ROS levels (**Fig. 1C**), and activation of pro-inflammatory signaling (**Figs. 1E and 1F**). Our results thus show that MNRR1 reduction results from stabilization of YME1L1 protease upon phosphorylation by ATM kinase.

### ROS generated by NOX2 activates ATM kinase in bacterial endotoxin treated placental cells *in vitro*

To probe in more detail the upstream basis of enhanced YME1L1 stability, we noted a previously defined inflammatory pathway in which activation of ATM kinase by NOX2 was demonstrated [48]. To determine whether this pathway was operating here we first examined whether NOX2 increased in LPS treated cells and found a robust increase (**Fig. 5A**). We then inhibited NOX2 to ask whether doing so prevented the LPS-dependent reduction in MNRR1 levels and found that MNRR1 was stabilized by the NOX2 inhibitor GSK2795039 (**Fig. 5B**).

**Figure 5:**
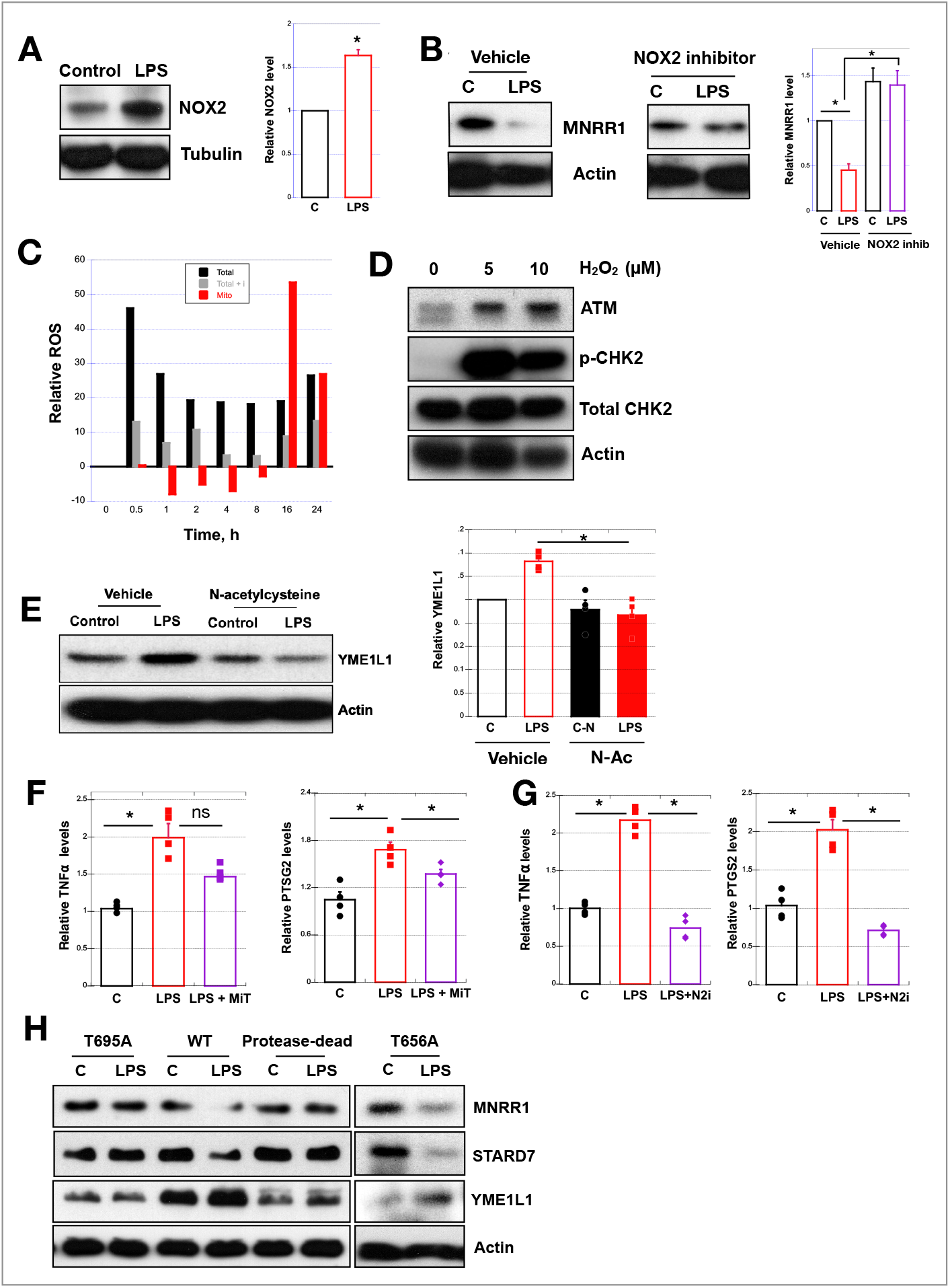
ROS generated by NOX2 activates ATM kinase in LPS treated placental cells. **(A)** *Left*, Equal amounts of HTR cells treated with control (water) or LPS were separated on an SDS-PAGE gel and probed for NOX2. Tubulin was probed as loading control. *Right*, NOX2 levels relative to tubulin are shown. **(B)** *Left*, Equal amounts of HTR cells were treated for 24 h with control (water) or LPS and, for 2^nd^ blot, 25 μM NOX2 inhibitor (using DMSO in control); lysates were separated on an SDS-PAGE gel and probed for NOX2. Actin was probed as loading control. *Right*, Relative MNRR1 levels are shown for each lane. **(C)** HTR cells were treated with control (water) or LPS for the times shown and ROS levels were measured as in Figure 1C. Total ROS, black; mitochondrial ROS, red; total ROS with ATM inhibitor, grey. **(D)** Equal amounts of HTR cells were treated with control (water) or hydrogen peroxide (H_2_O_2_) for 16 h and lysates separated on an SDS-PAGE gel and probed for p-CHK2, total CHK2, and ATM kinase. Actin was probed as loading control. **(E)** *Left*, Equal amounts of HTR cells treated with control (water) or LPS with either Vehicle (DMSO) or 100 μM N-acetyl cysteine for 24 h were separated on an SDS-PAGE gel and probed for YME1L1. Actin was probed as loading control. *Right*, Relative YME1L1 levels are shown for each condition. **(F)** *TNFα* and *PTGS2* transcript levels relative to *Actin* were measured in HTR cells treated with Control (water), LPS, or LPS + 20 μM MitoTempo. **(G)** *TNFα* and *PTGS2* relative transcript levels in HTR cells treated with Control (DMSO), LPS (LPS+DMSO) or LPS + 25 μM NOX2 inhibitor.

If a NOX2-generated “ROS burst” is upstream of mitochondrial ROS [49], we hypothesized that we should be able to detect this before a peak in mitochondrially-generated ROS. Indeed, following LPS treatment we saw that total ROS peaks within 30 minutes (black bars) whereas mitochondrial ROS (red bars) peaks at about 16 hours (**Fig. 5C**). Furthermore, the increase seen in total ROS was blocked with a NOX2 inhibitor (grey bars), suggesting that the ROS generated by NOX2 can activate ATM kinase (**Fig. 5C**). We tested ROS activation of ATM kinase in placental cells by generating ROS with hydrogen peroxide. We again saw an increase in ATM kinase amount as well as increased phosphorylation of a known ATM kinase target CHK2 [50] (**Fig. 5D**). We then tested the converse – whether scavenging ROS (with N-acetyl cysteine) would prevent LPS-stimulated phosphorylation of YME1L1 and found that such phosphorylation was indeed blocked (**Fig. 5E**). Taken together, these data suggest that ATM kinase is activated by NOX2-generated ROS and can phosphorylate and thereby stabilize YME1L1, which in turn reduces MNRR1 levels.

To assess whether ROS induced signaling was responsible for inflammation, we tested whether scavenging mitochondrial ROS or NOX2-mediated ROS would affect two markers of inflammation – *TNFα* (tumor necrosis factor-α) and *PTGS2* (prostaglandin synthase 2; also cyclooxygenase-2). We found that scavenging mitochondrial ROS (using MitoTempo, a mitochondria-specific ROS scavenger [51]) could partially reduce an LPS-induced increase in *TNFa* and *PTGS2* transcript levels (**Fig. 5F**). The use of the NOX2 inhibitor, on the other hand, could completely protect the increase in the same markers (**Fig. 5G**), suggesting that mitochondrial ROS is downstream of the NOX2-induced cytoplasmic ROS and that scavenging mitochondrial ROS only partially inhibits inflammation.

To further investigate YME1L1 phosphorylation, we generated a non-phosphorylatable point mutation (T695A) at the predicted target, threonine 695 (**Supplementary Figure 4A**). We tested the effect of this mutation in YME1L1-KO 293 cells by overexpressing this T695A mutant, WT, or PD-YME1L1. Doing so we found that the T695A mutation prevented the LPS-stimulated reduction in MNRR1 levels that is seen when the WT form is present (**Fig. 5H**), suggesting this phosphorylation is necessary for LPS-induced stabilization of YME1L1. Furthermore, this mutation behaves like PD-YME1L1 with respect to its known substrate STARD7 (**Fig. 5H**). The T695A mutation thus acts in a similar manner to PD-YME1L1. A second, control mutation, T656A, at a different threonine residue with a canonical ATM kinase recognition motif [52], does not prevent LPS-stimulated MNRR1 reduction (**Fig. 5H**), supporting the specificity of the T695 phosphorylation site in response to LPS treatment.

### Novel TLR4-independent signaling pathway is responsible for MNRR1-dependent reduction in mitochondrial function in bacterial endotoxin treated placental cells *in vitro*

Canonical LPS signaling is initiated by binding to Toll Like Receptor 4 (TLR4) [53–56]; therefore, we next asked if overexpression of TLR4 activates the NOX2-ATM-MNRR1 signaling pathway. We found that, although overexpression of TLR4 increases MNRR1 levels, LPS treatment reduces MNRR1 similarly to control cells (**Fig. 6A**). Since inflammation caused by LPS can occur either through MyD88-dependent signaling or MyD88-independent (TBK1-dependent) signaling [57–59] (**Fig. 6B**), we next assessed whether MNRR1 levels are reduced in mouse livers from WT or Myd88^−/−^ mice challenged with LPS. We found that MNRR1 levels are reduced in both WT and Myd88^−/−^ mice (**Fig. 6C**). Furthermore, examining activation of the kinase promoting the MyD88-independent immune response, we also found no change in TBK1 phosphorylation in HTR placental cells (**Supplementary Fig. 5A**). Taken together, these results eliminate the canonical TLR4 signaling pathway as the mediator of mitochondrial dysfunction.

**Figure 6:**
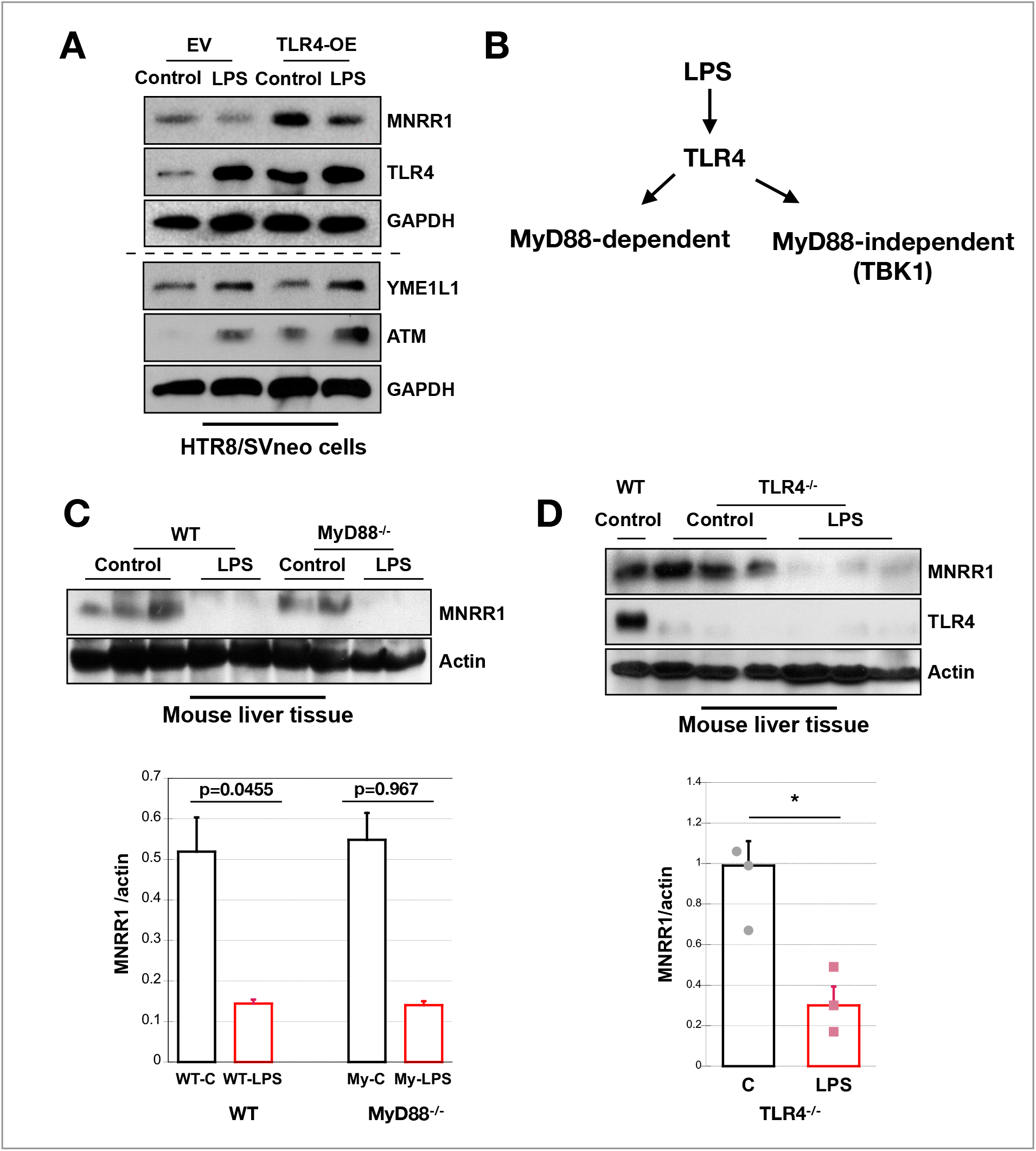
Novel TLR4-independent signaling pathway is responsible for MNRR1-dependent reduction in LPS treated placental cells. **(A)** Equal amounts of HTR cells overexpressing EV or TLR4 were treated or not with LPS, then lysates were separated on an SDS-PAGE gel and probed for MNRR1, TLR4, YME1L1, and ATM kinase. GAPDH was probed as loading control. **(B)** Schematic diagram for the two arms of the TLR4 signaling pathway. **(C)** *Above*, Equal amount of tissue lysates (WT or MyD88^−/−^) from mouse liver injected intraperitoneally with PBS (control) or LPS were separated on an SDS-PAGE gel and probed for MNRR1. Actin was probed as loading control. *Below*, The graph shows relative MNRR1 levels. **(D)** *Above*, Equal amount of liver lysates from mice (WT or TLR4^−/−^) that had been injected intraperitoneally with PBS (control) or LPS were separated on an SDS-PAGE gel and probed for MNRR1. Actin was probed as loading control. *Below*, Graph shows relative MNRR1 levels on blots.

We then hypothesized that TLR4 may directly interact with NOX2 to initiate this pathway, and hence tested MNRR1 levels in TLR4^−/−^ mouse liver tissue lysates injected with PBS (control) or LPS. We found that MNRR1 levels are reduced also in TLR4^−/−^ mouse livers challenged with LPS (**Fig. 6D**). Besides MNRR1, we tested for other markers (NOX2 and ATM kinase) both in the LPS-injected mouse placentas (**Supplementary Fig. 5B**), where we originally found a reduction in MNRR1 (**Fig. 1A**), as well as in the TLR4^−/−^ mouse liver tissue lysate (**Supplementary Fig. 5C**). We found the pathway to be active even in the animal samples, consistent with the results found in the human cell culture system (**Supplementary figure 4B and Fig. 5A**), confirming that the same pathway is active is the TLR4^−/−^ animals. To verify that the TLR4-independent reduction in MNRR1 levels seen in the TLR4^−/−^ animals is initiated by NOX2 activation, we used a NOX2 inhibitor in TLR4^−/−^ mouse macrophages [60] and found that the NOX2 inhibitor prevents LPS-induced reduction in MNRR1 (**Supplementary Fig. 5D**). Taken together, we conclude that LPS acts through a TLR4-independent pathway to activate ATM kinase to phosphorylate YME1L1 at Thr-656, stabilizing it and thereby reducing MNRR1 levels.

### MNRR1 functions as an anti-inflammatory effector via its nuclear function

To confirm that MNRR1 is upstream of the inflammatory signaling we generated a MNRR1-depleted human placental cell line and assessed transcript levels of two inflammatory markers – phospho-JNK (**Supplementary Fig. 5E**) and *TNFα* (**Supplementary Fig. 5F**) – and found these to be increased. Since MNRR1 is present in both the nucleus and the mitochondria and has a different function in each [27], we investigated the compartment-specific effect of LPS on MNRR1 by assessing OCR and the stimulation of inflammation-associated genes *TNFα* and *PTGS2*. We found that LPS treatment increased the transcript levels of both these genes and that overexpression of either WT or the C-S mutant of MNRR1 that does not localize to mitochondria [27] can prevent this increase (**Figs. 7A, 7B**), as can the MNRR1 activator Nitazoxanide (N) (**Figs. 7C, 7D**). To determine whether the anti-inflammatory role is due to nuclear function of MNRR1, we asked if overexpressing CHCHD4, which is required for MNRR1 import into the mitochondria [27], could prevent the LPS-induced deficit in oxygen consumption. We found that CHCHD4 overexpression can increase oxygen consumption (**Supplementary Fig. 5G**), as also shown previously [61], but that LPS-treatment reduces oxygen consumption to the same extent as seen in the absence of CHCHD4 overexpression, suggesting that specifically nuclear MNRR1 is required to prevent inflammation.

**Figure 7:**
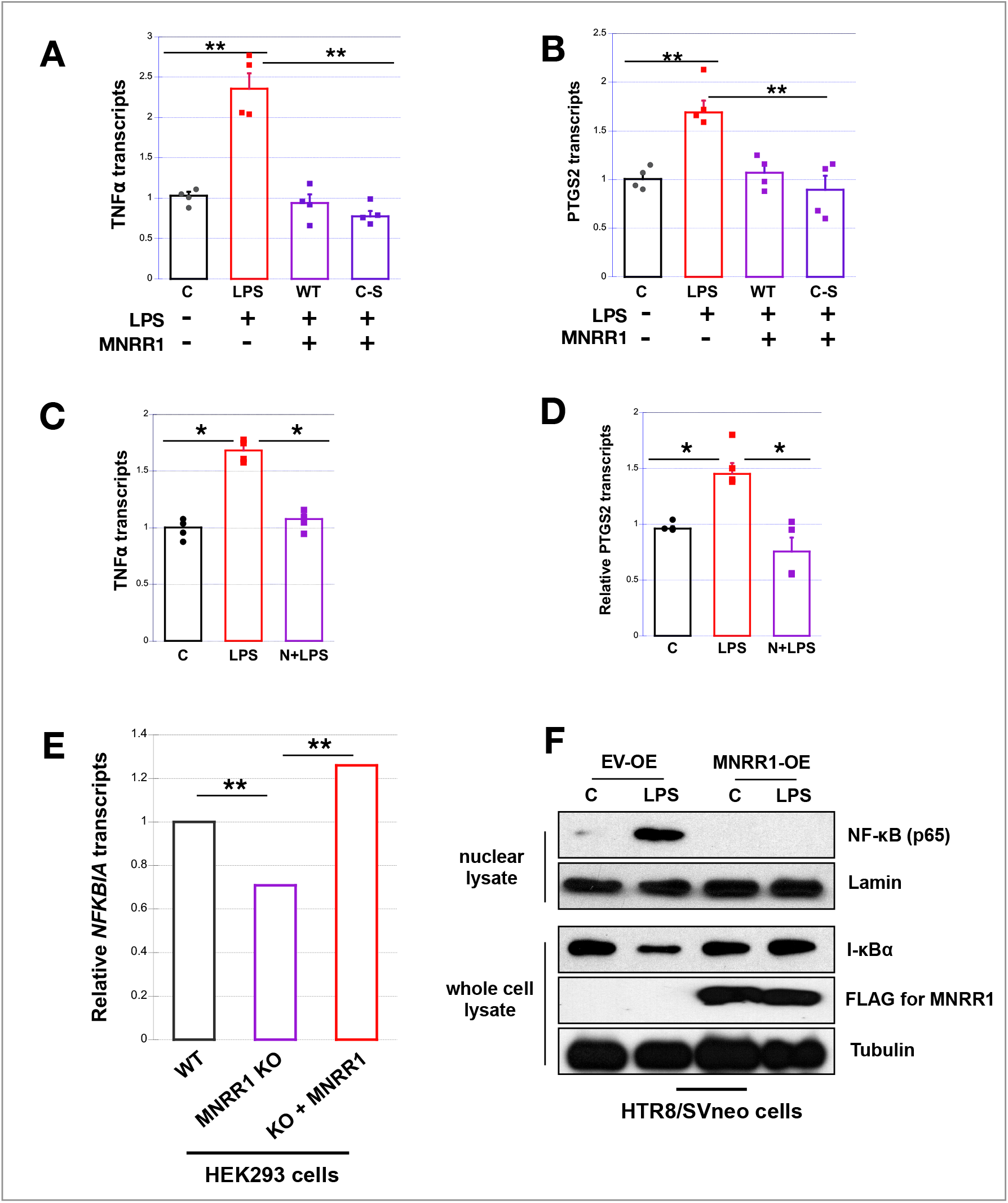
MNRR1 functions as anti-inflammatory via its nuclear function. Relative *TNFα* **(A)** and *PTGS2* **(B)** transcript levels in HTR cells treated with Control (EV), LPS (EV + LPS), WT-MNRR1 (WT + LPS), or C-S-MNRR1 (C-S + LPS). **(C)** Relative *TNFα* and **(D)** *PTGS2* transcript levels in HTR cells treated with Control (DMSO), LPS (LPS + DMSO), or LPS + 10 μM Nitazoxanide. **(E)** RNA-sequencing (HEK293 cells) showing that *NFKBIA* transcript levels are significantly reduced in MNRR1 knockout cells (KO) relative to wild type controls (WT). This reduction is rescued by overexpressing the transcriptionally active mutant of MNRR1 (K119R-MNRR1). **(F)** Nuclear NF-κB EV or MNRR1 and treated with control or LPS. Lamin was probed as a nuclear loading control. Whole cell lysates from the same experiment were probed for I-κB and FLAG (MNRR1) levels. Tubulin was probed as loading control.

To probe the mechanism by which nuclear MNRR1 can inhibit inflammation, we looked to see if any regulatory components of the NF-κB signaling pathway are transcriptionally regulated by MNRR1. We found that I-κBα *(NFKBIA)*, a regulator that binds NF-κB and retains it in the cytoplasm [62], is transcriptionally activated by MNRR1 (**Fig. 7E**). Consistent with the prediction from RNA-sequencing, performed in HEK293 cells [63], we found in the placental cells that I-κBα levels are reduced by LPS treatment (**Fig. 7F**), thereby allowing nuclear localization of NF-κB. Overexpression of MNRR1, however, can prevent these effects (**Fig. 7F**). These findings suggest that MNRR1 can act as an anti-inflammatory agent at least in part by preventing activation of NF-κB. Since one of the classic targets of NF-κB, *COX-2*, is required for induction of labor under physiological conditions [64, 65], MNRR1 expression may blunt the effects of inflammation by preventing nuclear translocation of NF-κB.

## Discussion

Acute inflammation due to intra-amniotic infection is causally linked to spontaneous premature labor [66–69]. Mitochondria can serve as an early sensor of inflammatory stress [70–74]. Here, we show that MNRR1 is reduced by a post-translational mechanism in *in vivo* and *in vitro* models of placental inflammation, leading to the generation of mitochondrial ROS. Surprisingly, the mitochondrial ROS that is the source of inflammatory signaling in the placenta takes place via a TLR4-independent signaling pathway. This pathway is initiated by activation of ATM kinase via NOX2-dependent ROS, which in turn phosphorylates YME1L1, a protease that degrades MNRR1 in the mitochondria. Although ATM kinase is primarily activated in response to DNA damage [75], it was also shown be activated in response to LPS treatment in macrophage cells [76] although the downstream signaling targets were not identified. A recent study in a renal tubular epithelial cell model of LPS-induced sepsis [77] also identifies ATM activation as playing a key role in inflammation and autophagy activation. ATM kinase is localized to mitochondria [78], more specifically to the inner mitochondrial membrane [79], the same sub-mitochondrial compartment that harbors YME1L1 [80, 81]. YME1L1, although embedded in the inner membrane, exposes a large catalytic domain facing the IMS [80]. We showed that ATM-mediated threonine phosphorylation of YME1L1 can enhance its effective activity, resulting in faster turnover of MNRR1 in the mitochondria. Depletion of MNRR1 results in reduced oxygen consumption, reduced ATP levels, and increased ROS. These changes activate an irreversible and self-amplifying inflammatory signaling cascade that may disrupt signaling at the fetal-maternal interface.

MNRR1 has previously been associated with several diseases both in terms of altered expression and through mutations. Depleted MNRR1 protein levels have been found in an *in vivo* model for juvenile Niemann Pick type C disease [82] and *in vitro* model for MELAS (Mitochondrial Encephalomyopathy Lactic Acidosis and Stroke-like episodes) syndrome [32]. Mutations in MNRR1 have been associated with a number of neurodegenerative diseases such as Parkinson’s [83–85], Alzheimer’s [86, 87], and Charcot-Marie-Tooth disease type 1A [28]. Of note, MELAS, caused by a mtDNA mutation in the mitochondrial tRNA^Leu(UUR)^ gene (m.3243A > G), has been associated with spontaneous preterm birth [88, 89] and increased incidence of preeclampsia and gestational diabetes mellitus [88]. We have recently shown by overexpression of MNRR1 that oxygen consumption and other deficits associated with MELAS can be rescued in a cell culture model for the disease [32]. MNRR1 activation, either by overexpression or, more interestingly, by using a chemical activator, can thus provide multiple benefits that protect placental mitochondria and reduce inflammation.

MNRR1 is known to function in both the nucleus and the mitochondria [26–29] and as a nuclear transactivator can promote its own transcription [27]. Although we have focused here on the consequences of mitochondrial depletion, the rescue by pharmacological activation of transcription (**Fig. 1F**) and by overexpression of MNRR1 that cannot enter the mitochondria (**Figs. 7A, 7B**) suggests that activating its nuclear function could suffice to prevent placental damage. In addition to activating itself, MNRR1 is a transcriptional activator for ROS scavenging enzymes such as *SOD2* (superoxide dismutase) and *GPX* (glutathione peroxidase) [27] and also is a regulator of mitophagy genes such as *ATG7* and *PARK2* (Parkin) [32]. A similar conclusion about the importance of its nuclear function was reached in a MELAS model [32]. Besides MNRR1’s ability to regulate genes involved in ROS scavenging, we now identify another transcriptional target – IκBα – that can contribute to its anti-inflammatory role via inhibition of NF-κB.

The novel NOX2/ATM kinase/YME1L1/MNRR1/COX-2 axis we have described provides insight into the mechanism by which placental inflammation can lead to preterm labor and birth (**Fig. 8**). There are multiple points at which we could modulate this pathway but activation of MNRR1 may be an ideal point to break the cycle of ROS-induced inflammation. COX-2 was initially considered an ideal target since pharmacological inhibition of COX-2 can prevent inflammation induced preterm labor [90] in mice. However, an offsetting consideration is that COX-2 is important in physiological labor and its loss can impair closure of the ductus arteriosus [91, 92]. This temporally defined role of COX-2 [93] has led to concerns about using cyclooxygenase inhibitors in the clinic during pregnancy [94]. Another current treatment uses steroidal compounds in the antenatal period to prevent respiratory distress syndrome and mortality in anticipated cases of preterm birth [95]. However, steroids are not always useful and have been associated with deleterious effects both on the fetus such as cerebral palsy [96], microcephaly [97–99], lower birthweight [100–102], adrenal suppression [103], the development of impaired glucose tolerance and hypertension later in life [97, 104], and on the mother such as risk of infection [99], loss of glycemic control in diabetics [105, 106], suppression of the hypothalamic axis [107], and reduced fetal growth velocity [108]. Since the pro-inflammatory signaling proceeds through degradation of mitochondrial MNRR1 whereas its nuclear function is sufficient to rescue the effects of LPS, transcriptional activation of MNRR1 can provide a treatment option. In addition, the recent demonstration that MNRR1 activation may be able to augment or in some cases even replace steroids for respiratory distress syndrome [109] adds additional impetus for development of this targeted therapy.

**Figure 8:**
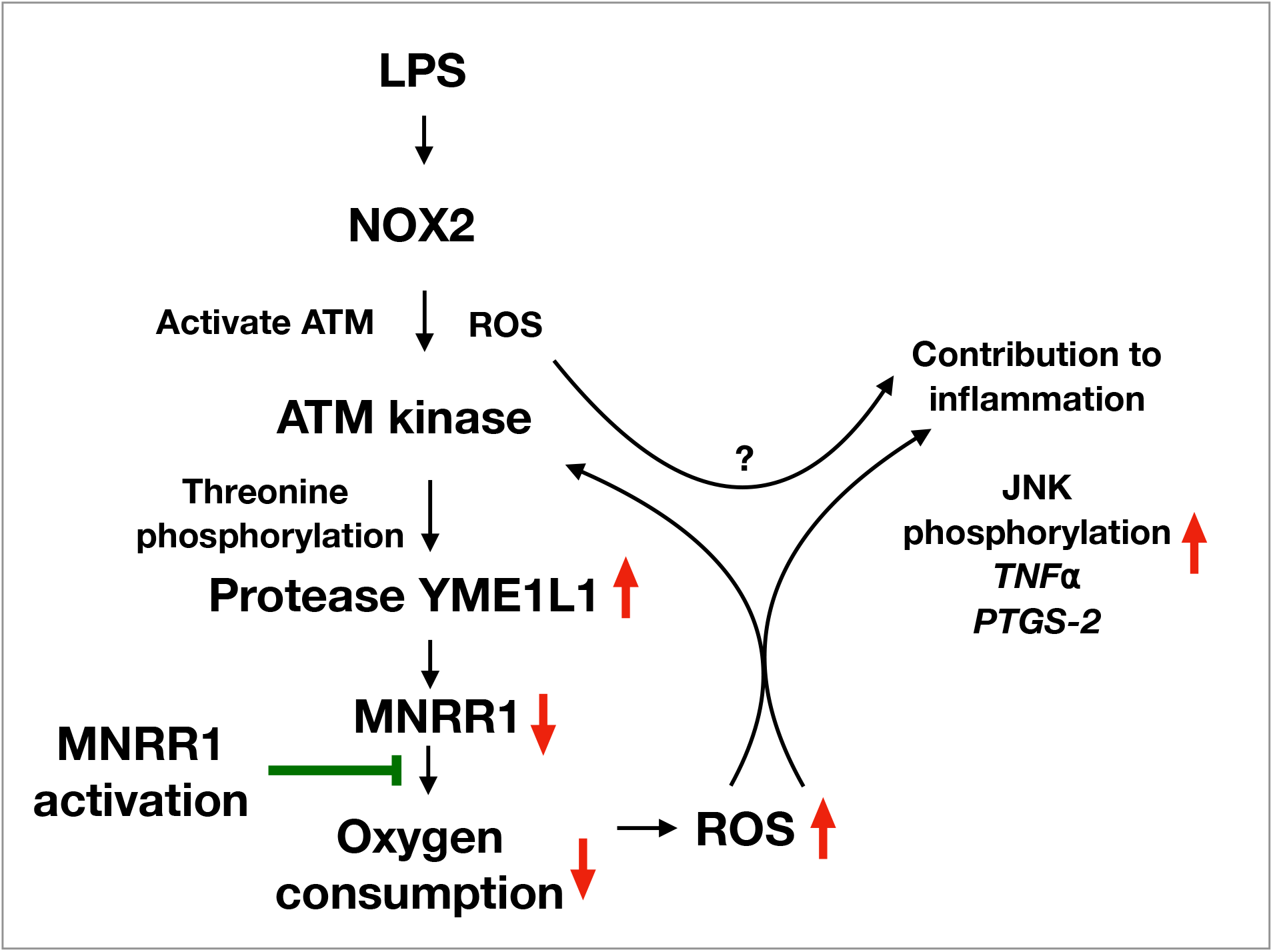
Model of MNRR1 action to suppress inflammation. Schematic summary of the role of MNRR1 in inducing inflammation. Bacterial endotoxin activates ATM kinase via NOX2-mediated ROS. Increased ATM activity in turn stabilizes YME1L1 protease by enhancing its threonine phosphorylation. Increased YME1L1 protease degrades MNRR1 to reduce oxygen consumption and increase ROS levels that contribute towards an inflammatory phenotype as evidenced by increased levels of *TNFα* and *cyclooxygenase-2* (*PTGS2*). Activation of MNRR1 can prevent a reduction in mitochondrial function and increase of ROS levels, thereby preventing inflammation.

In summary, we have identified a novel signaling axis by which inflammation induced by the bacterial mimetic LPS causes mitochondrial dysfunction. It does so by reducing the level of the bi-organellar regulator MNRR1 in response to phosphorylation and stabilization of IMS protease YME1L1, which turns over MNRR1. Phosphorylation is carried out by ATM kinase after activation by NOX2-produced ROS promoted by LPS. The mitochondrial ROS that stem from MNRR1 inhibition cause JNK phosphorylation and consequent activation of the cytokines TNFα and cyclooxygenase 2.

## Experimental Procedures

### Cell culture and reagents

#### Cell lines

All cell media were supplemented with 10% fetal bovine serum (FBS) (Sigma Aldrich, St. Louis, MO) plus Penicillin Streptomycin (HyClone, Logan, UT). HTR8/SVNeo (HTR), RAW, and JAR cells were cultured in Roswell Park Memorial Institute Medium (RPMI) (HyClone, Logan, UT). The BeWO cells were grown in F12K media (Gibco, Waltham, MA). HEK293 cells were grown in Dulbecco’s modified Eagle’s medium (without pyruvate). The WT and TLR4^−/−^ immortalized macrophages and HepG2 cells, were grown in Dulbecco’s modified Eagle’s medium (with 1 mM pyruvate). YME1L1 knockout-HEK293 cells were grown in Dulbecco’s modified Eagle’s medium (with 1 mM pyruvate) supplemented with non-essential amino acids (Gibco).

#### Chemicals

Nitazoxanide and Clotrimazole were obtained from Selleckchem (Houston, TX) and solubilized in DMSO (used as vehicle control in all experiments with these compounds). Ultrapure LPS for cell culture experiments (Lipopolysaccharide from *Escherichia coli* 0111:B4) was purchased from Invivogen. NOX2 inhibitor GSK2795039 was obtained from MedChem Express (Monmouth Junction, NJ). ATM inhibitor KU-55933 was from Cell Signaling Technology.

#### Plasmids

The WT and protease dead (PD) YME1L1 plasmids were a kind gift from Dr. Thomas Langer, University of Cologne, DE. The *MNRR1* promoter luciferase reporter plasmid has been described previously [27]. The human TLR4 overexpression plasmid was purchased from Addgene (Cat. #13086). The T695A and T656A mutations were generated in WT-YME1L1 plasmid via QuikChange Lightning Site-Directed Mutagenesis Kit (Agilent, Santa Clara, CA) and confirmed by sequencing. All the expression plasmids were purified using the EndoFree plasmid purification kit from Qiagen (Valencia, CA).

### Transient transfection of HTR cells

HTR cells were transfected with the indicated plasmids using TransFast transfection reagent (Promega, Madison, WI) according to the manufacturer’s protocol. A TransFast:DNA ratio of 3:1 in serum and antibiotic free medium was used. Following incubation at room temperature for ~15 min, the cells were overlaid with the mixture. The plates were incubated for 1 h at 37 °C followed by replacement with complete medium and further incubation for the indicated time.

### Real-time polymerase chain reaction

Total cellular RNA was extracted from mouse placental tissue or HTR cells with an RNeasy Plus Mini Kit (Qiagen, Valencia, CA) according to the manufacturer’s instructions. Complementary DNA (cDNA) was generated by reverse transcriptase polymerase chain reaction (PCR) using the ProtoScript^®^ II First Strand cDNA Synthesis Kit (NEB, Ipswich, MA). Transcript levels were measured by real time PCR using SYBR green on an ABI 7500 system. Real-time analysis was performed by the ΔΔ^Ct^ method. The primer sequences used were as follows (F, forward; R, reverse): Mouse MNRR1: F- ATGGCCCAGATGGCTACC, R- CTGGTTCTGAGCACACTCCA; Mouse Actin: F- TCCTCCCTGGAGAAGAGCTA, R- ACGGATGTCAACGTCACACT; Human MNRR1: F- CACACATTGGGTCACGCCATTACT, R- TTCTGGGCACACTCCAGAAACTGT; Human Actin: F- CATTAAGGAGAAGCTGTGCT, R- GTTGAAGGTAGTTTCGTGGA; Human YME1L1: F- TGAAGGGGTTTCTTTTGCGG, R- TCGCCTTAGGGAATCATTGGT, Human TNFα-F- TGTAGCAAACCCTCAAGCTG, R- GAGGTTGACCTTGGTCTGGT, Human PTGS2 F-ATGATGTTTGCATTCTTTGCCCAG, R- CATCCTTGAAAAGGCGCAGTTTA

### Luciferase reporter assay

Luciferase assays were performed with the dual-luciferase reporter assay kit (Promega, Madison, WI) per the manufacturer’s instructions. Transfection efficiency was normalized with the co-transfected pRL-SV40 *Renilla* luciferase expression plasmid [26, 28, 29].

### Intact cellular oxygen consumption

Cellular oxygen consumption was measured with a Seahorse XF^e^24 Bioanalyzer (Agilent, Santa Clara, CA). Cells were plated at a concentration of 3 × 10^4^ per well a day prior to treatment and basal oxygen consumption was measured 24 h after treatments as described [28, 29].

### ATP measurements

7.5 × 10^4^ HTR cells per well were plated on a 96-well plate a day prior to treatment and ATP levels were measured using Cell Titer Glo (Promega, Madison, WI) according to manufacturer’s instructions.

### ROS measurements

Total cellular ROS measurements were performed with CM-H2DCFDA (Life Technologies, Grand Island, NY). Cells were distributed into 96-well plates at 7.5 × 10^4^ cells per well and incubated for 24 h or as described in specific experiments. Cells were then treated with 10 μM CM-H_2_DCFDA in serum- and antibiotic-free medium for 1 h. Cells were washed twice in phosphate buffered saline and analyzed for fluorescence on a Gen5 Microplate Reader (BioTek Inc, Winooski, VT). For mitochondrial ROS measurements, the cells were treated as above but with 5 μM Mitosox Red (Life Technologies) for 30 min.

### Confocal microscopy

Confocal microscopy was performed as described [28, 29]. For mouse placental tissue sections, 8-10 μm thick transverse sections were stained with anti-MNRR1 antibody (1:50 Proteintech Inc., Chicago, IL). The secondary antibody used was donkey anti-rabbit IgG Alexa 594 (1:200, Jackson Labs, Bar Harbor, ME). These were imaged with a Leica TCS SP5 microscope and images were combined in Photoshop. Co-localization (overlap of the two fluorophores) and intensity (number of pixels per unit area) were quantitated using Volocity image analysis software (Perkin Elmer, Waltham, MA). For human placental cells, staining was performed with anti-MNRR1 (1:50 Proteintech Inc., Chicago, IL) conjugated to CoraLite-594 and anti-TOM20 conjugated to CoraLite-488 (1:200, Proteintech Inc., Chicago, IL).

### Mitochondria isolation

Mitochondria were isolated from cells with a Mitochondrial Isolation Kit (Thermo Scientific, Rockford, IL) as described previously [28, 29].The nuclear fraction was obtained by low-speed centrifugation and the mitochondrial fraction was obtained after high-speed centrifugation of the nuclear supernatant. Cross-contamination between the fractions was analyzed with compartment-specific antibodies.

### Immunoblotting and co-immunoprecipitation

Immunoblotting on a PVDF membrane was performed as described previously [26, 27]. Unless specified otherwise, primary antibodies were used at a concentration of 1:500 and secondary antibodies at a concentration of 1:5000. The MNRR1 (19424-1-AP), YME1L1 (11510-1-AP), DRBP76 (19887-1-AP), MTCO2 (55070-1-AP), STARD7 (156890-1-AP), NOX2 (19013-1-AP), RelA/p65 (10745-1-AP), and TLR4 (19811-1-AP) (used for mouse tissue) antibodies were purchased from Proteintech Inc. (Chicago, IL). The GAPDH (8884), phospho-JNK (4668), total JNK (9252), anti-phosphothreonine (9386), ATM (2873), CHK2 (6334), phosphothreonine 68-CHK2 (2197), I-κBα (4814), phospho-TBK1 (5483), and total TBK1 (3504) antibodies were purchased from Cell Signaling Technology Inc (Beverly, MA). The Lamin (sc-6217) and TLR4 (sc-293072) antibodies (used for human cell lines) were purchased from Santa Cruz Biotechnology (SCBT), Inc, Dallas, TX). The FLAG antibody (A8592) was purchased from Sigma. Co-immunoprecipitation experiments were performed according to the supplier’s protocol by incubating the antibody-adsorbed beads overnight at 4 °C. For co-immunoprecipitation, phosphothreonine antibody (Cell Signaling) conjugated to L-agarose beads (sc-2336, SCBT) or YME1L1 antibody (Proteintech) conjugated to protein A/G-agarose beads (sc-2003, SCBT) were used.

### Statistical analysis

Statistical analyses were performed with MSTAT version 6.1.1 (N. Drinkwater, University of Wisconsin, Madison, WI). The two-sided Wilcoxon rank-sum test was applied to determine statistical significance for *p*-values. Data were considered statistically significant with p <0.05. Error bars represent standard error of mean.

### Animal experiments and injections

Mouse placental samples: Samples were obtained using the intraperitoneal injection LPS (*Escherichia coli* O111:B4; Sigma) model that results in 100% preterm labor/birth [40]. Briefly, pregnant B6 mice were intraperitoneally injected on 16.5 dpc with 15 μg of LPS in 200 μL of PBS using a 26-gauge needle. Controls were injected with 200 μL of PBS. Mice were monitored via video recording using an infrared camera to determine gestational age and the rate of preterm labor. Placentas were collected before preterm birth (12-13 h after LPS injection).

Mouse liver samples for Myd88^−/−^ and TLR4^−/−^: Mice of approximately 3-months were injected intraperitoneally with LPS (*Escherichia coli* O111:B4, Sigma, 2 μg/gm body weight) or PBS for 18 h [110]. Liver tissues collected from the mice after LPS treatment were homogenized in NP-40 lysis buffer in the presence of proteasome inhibitors. Lipid contents were briefly extracted from the liver tissue lysates by the SDS buffer, and the protein supernatants were denatured for Western blot analyses [111].

All animal procedures were approved by the Institutional Animal Care and Use Committee (IACUC) at Wayne State University under Protocol No. A-07-03-15.

## Materials and data availability

The reagents and data from the current study are available from the corresponding authors on reasonable request.

## Acknowledgements

We thank Dr. Thomas Langer (University of Cologne, Cologne, Germany) for providing the YME1L1 plasmids, Dr. Kezhong Zhang (Wayne State University, Detroit, MI, US) for providing MyD88^−/−^ tissues and TLR4^−/−^ tissue lysates, and Dr. Douglas Golenbock (University of Massachusetts Medical School, Worcester, MA, US) for providing the TLR4^+/+^ and TLR4^−/−^ mouse macrophages. This work was supported by the Perinatology Research Branch, Division of Obstetrics and Maternal-Fetal Medicine, Division of Intramural Research, *Eunice Kennedy Shriver* National Institute of Child Health and Human Development, National Institutes of Health, U.S. Department of Health and Human Services (NICHD/NIH/DHHS); and, in part, with Federal funds from NICHD/NIH/DHHS under Contract No. HHSN275201300006C. Dr. Romero has contributed to this work as part of his official duties as an employee of the United States Federal Government. Opinions, interpretations, conclusions, and recommendations are those of the authors and are not necessarily endorsed by the NIH.

## Author contributions

NP performed all experiments, YK helped with generation of Western blots, NGL provided the human and mouse placental tissue samples. NP, SA, and LIG analyzed the results and participated in experimental design. YX, AF performed the screen to identify MNRR1 activators and inhibitors. NP and LIG wrote the manuscript. All authors reviewed the manuscript.

## Figure legends

**Supplementary Figure 1.**
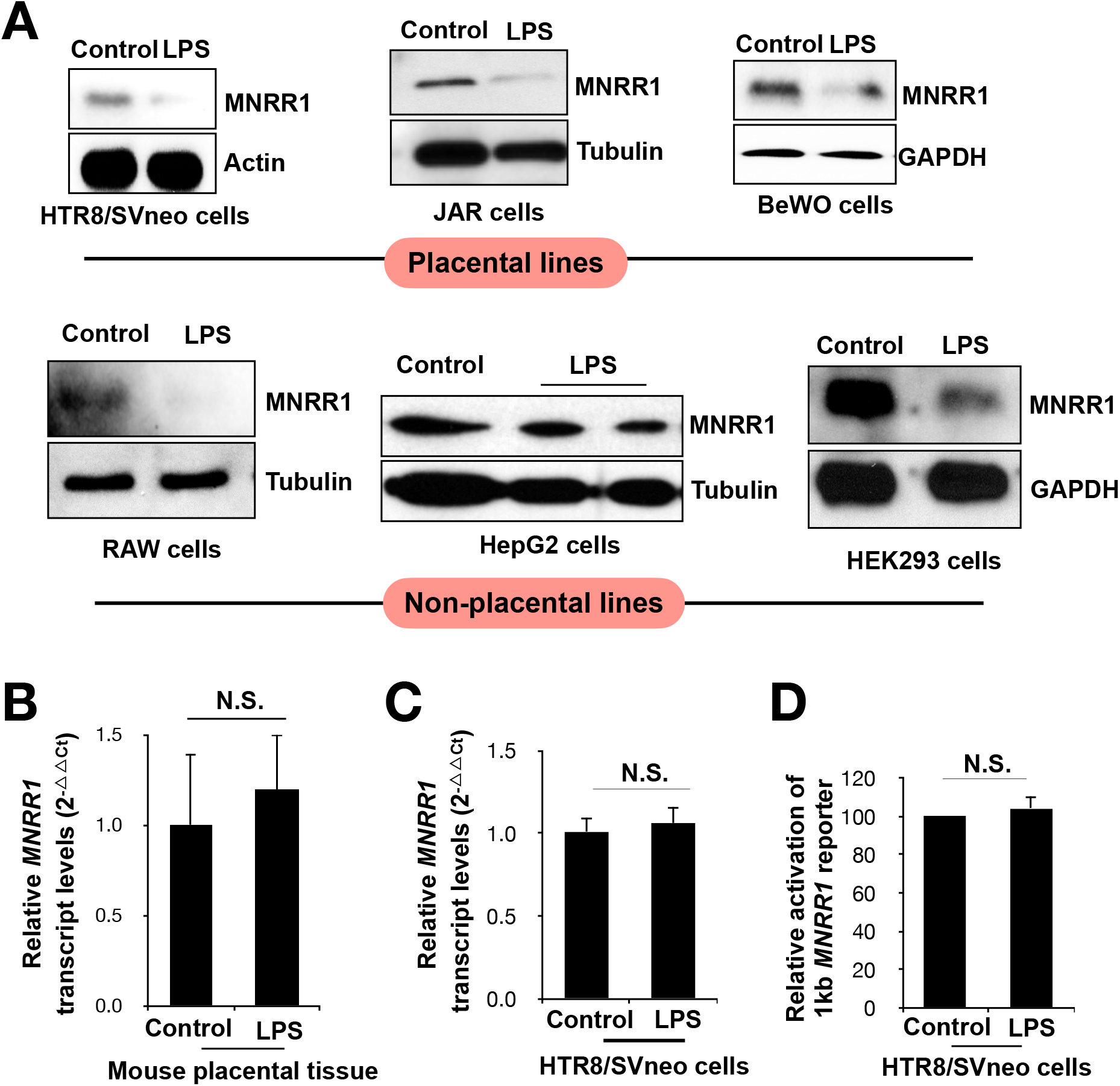
**(A)** Generality of LPS-stimulated reduction of MNRR1 levels by western analysis of lysates of various cell lines treated with control or LPS, then probed for MNRR1 and labeled loading control. *MNRR1* transcript levels are shown relative to *Actin* in mouse placental tissues **(B)**, and HTR cells **(C). (D)** Transcript levels of *MNRR1*-luciferase reporter in HTR cells.

**Supplementary Figure 2.**
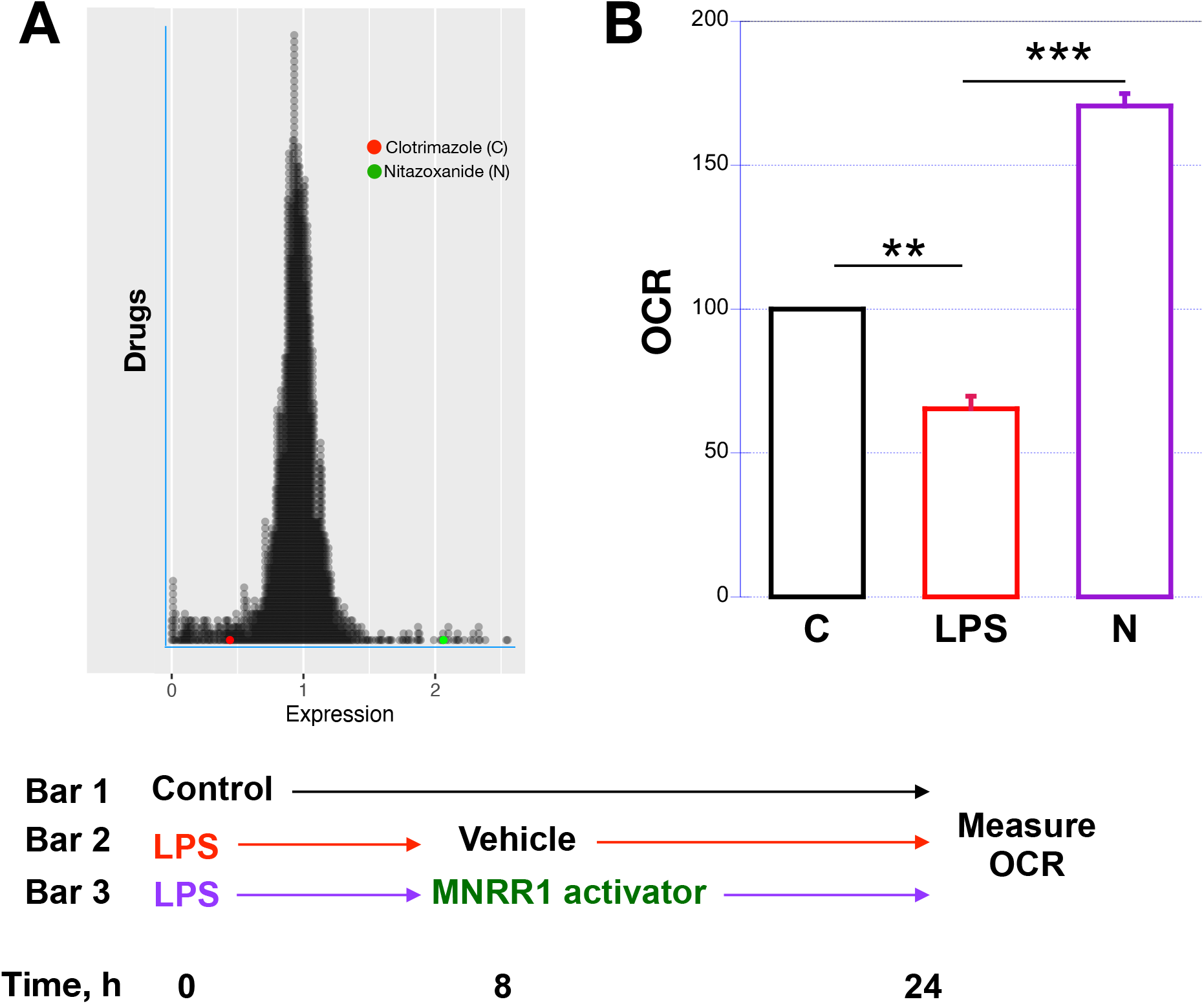
**(A)** Results of screen of 2400 FDA-approved drugs identified to transcriptionally activate (>1), inhibit (<1) or not affect (=1) MNRR1. Each circle represents one drug and the MNRR1 activator (Nitazoxanide (N), green) and MNRR1 inhibitor (Clotrimazole (C), red) have been highlighted. **(B)** Intact cellular oxygen consumption in HTR cells treated as described in the scheme below.

**Supplementary Figure 3.**
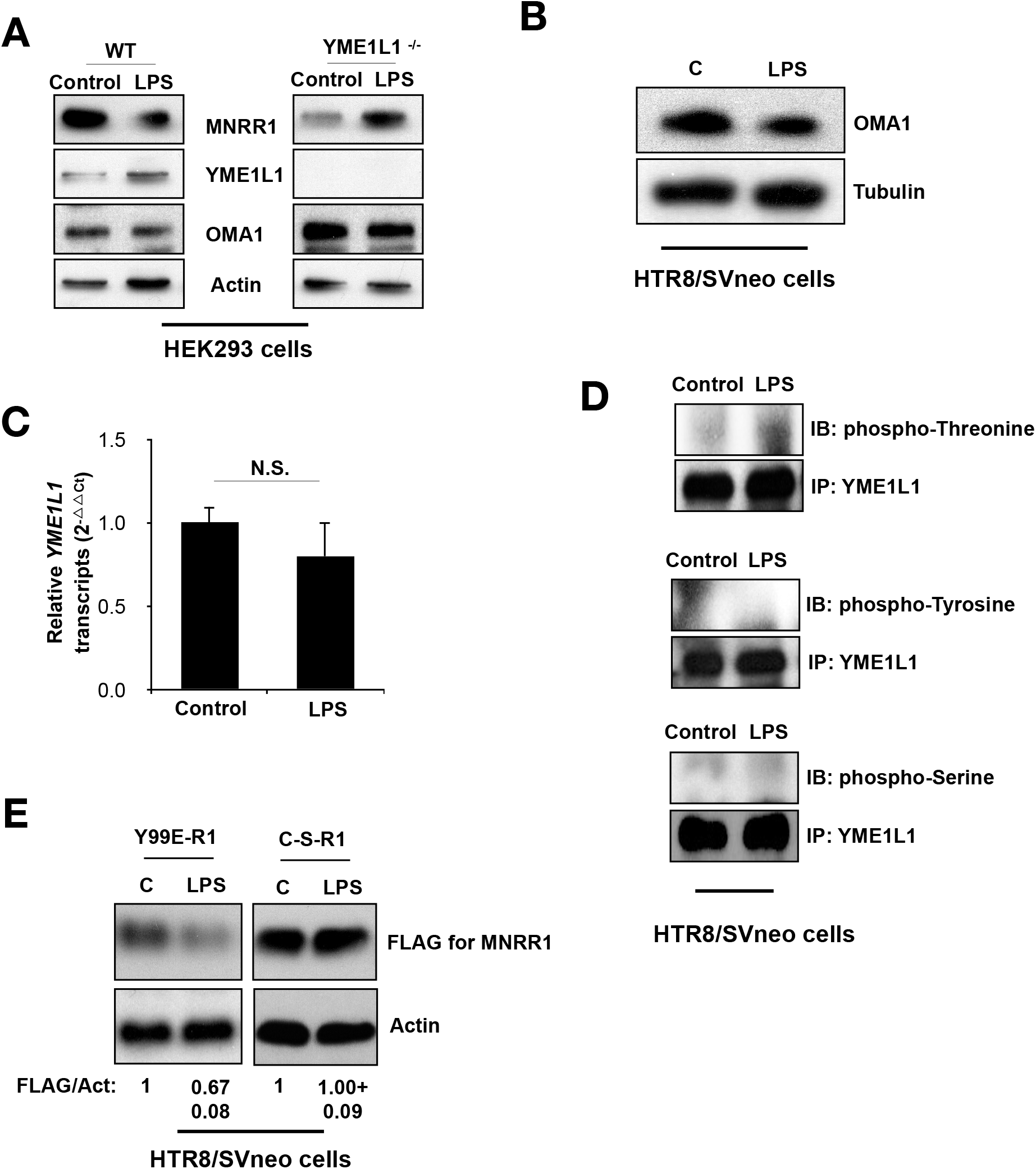
**(A)** Equal amounts of WT or YME1L1^−/−^ cells treated with control or LPS (1 μg/mL) for 24 h were separated on an SDS-PAGE gel and probed for MNRR1, YME1L1, and OMA1. Actin was probed as loading control. **(B)** Equal amounts of HTR cells treated with control (water) or LPS (500 ng/mL) were separated on an SDS-PAGE gel and probed for OMA1. Actin was probed as loading control. **(C)** *YME1L1* transcript levels relative to *Actin* in HTR cells. **(D)** HTR cells were treated with control (water) or LPS. Equal amounts of whole cell lysates were used for immunoprecipitation with a YME1L1 antibody. Equal amounts IP eluates were probed for p-Serine, p-Threonine, or p-Tyrosine and YME1L1 was probed to assess loading. **(E)** Equal amounts of HTR cells overexpressing FLAG-tagged Y99E or C-S-MNRR1 were treated with control or LPS and lysates were separated on an SDS-PAGE gel and probed for FLAG. Actin was probed as loading control.

**Supplementary Figure 4.**
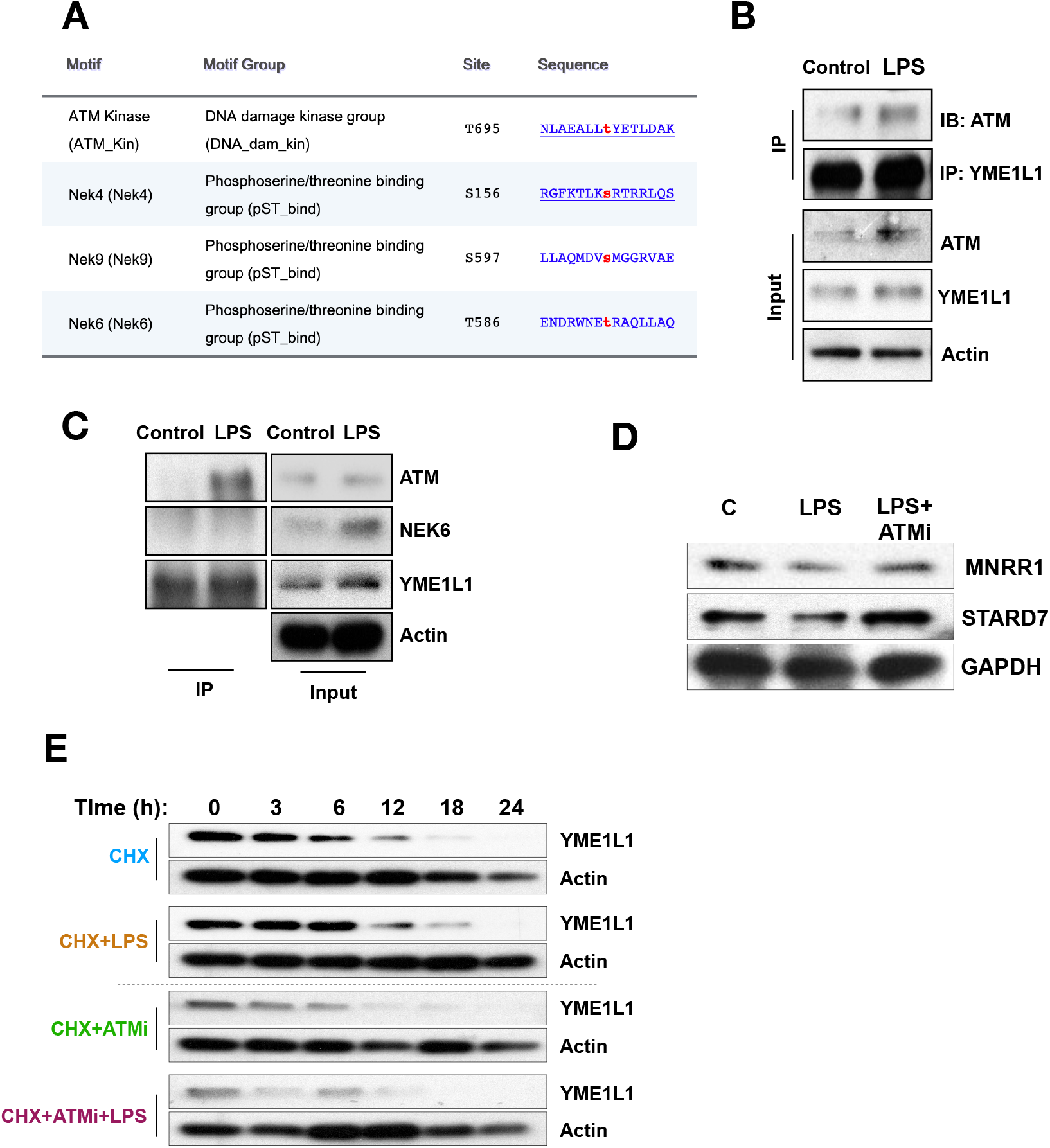
**(A)** Bioinformatic prediction from Scansite for kinases that might phosphorylate YME1L1. **(B)** Interaction of YME1L1 and ATM kinase. HTR cells were treated with water or LPS. Equal amounts of whole cell lysates were used for immunoprecipitation with YME1L1 antibody. Equal amounts IP eluates were probed for ATM kinase and YME1L1, the latter to assess loading. Input lysates were also probed for ATM and YME1L1 and Actin was probed as loading control. **(C)** Interaction of YME1L1 and NEK6 kinase. HTR cells were treated with water or LPS. Equal amounts of whole cell lysates were used for immunoprecipitation using YME1L1 antibody. Equal amounts IP eluates were probed for ATM kinase, NEK6, and YME1L1 was probed to assess loading. Input lysates were also probed for ATM, NEK6, and YME1L1 and actin was probed as loading control. **(D)** HTR cells were treated with vehicle, LPS, or LPS plus ATM inhibitor (1 μM) for 24 h. Equal amounts of cell lysates were separated on an SDS-PAGE gel and probed for MNRR1 and STARD7. GAPDH was probed as loading control. **(E)** Data used for time course of YME1L1 turnover (Figure 4D). Equal numbers of HTR cells were plated on a 6-well plate and treated with water or LPS and 100 μg/mL cycloheximide for the times shown. Equal amounts cell lysates were separated on an SDS-PAGE gel and probed for YME1L1. Actin was probed as loading control.

**Supplementary Figure 5.**
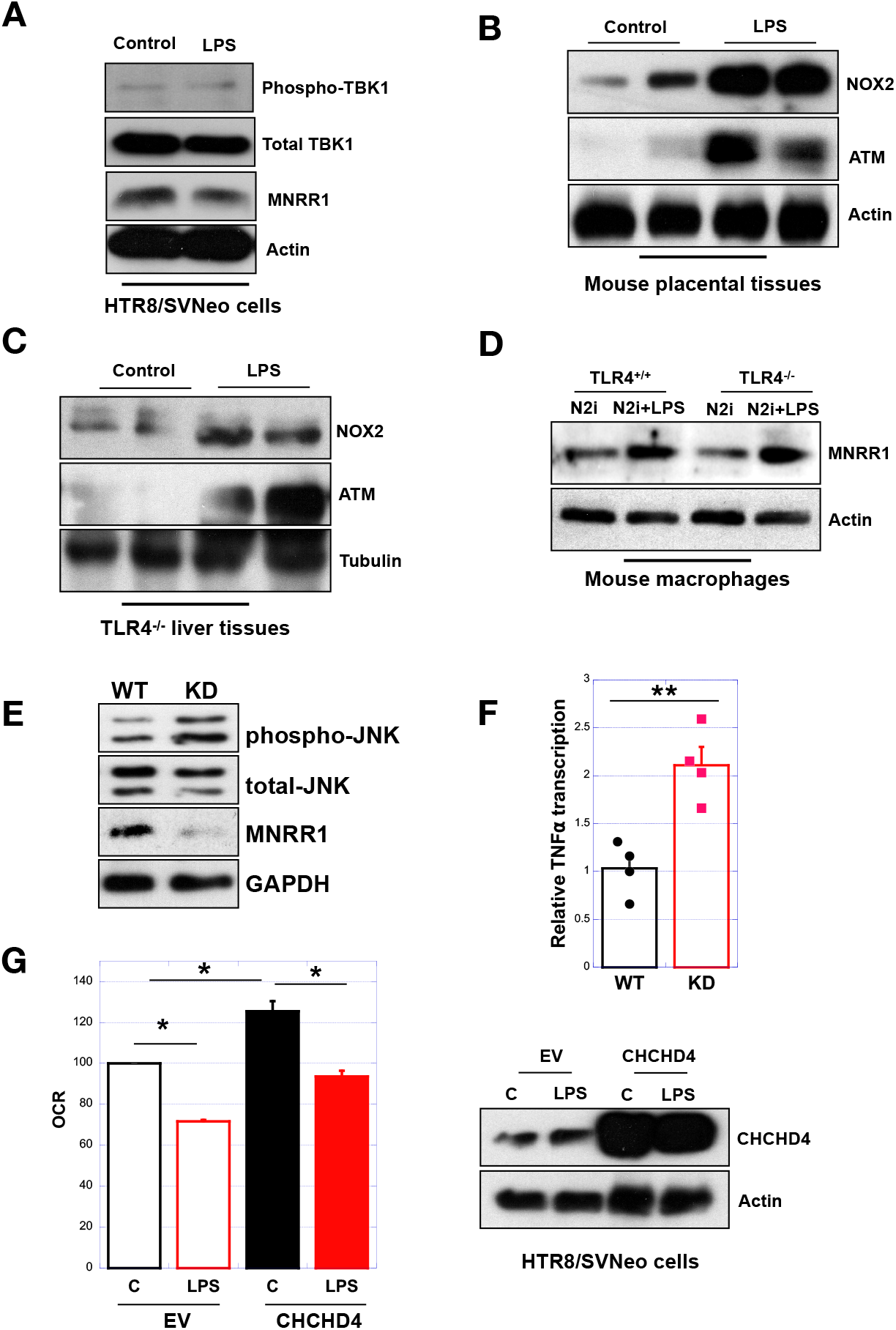
**(A)** Equal amounts of HTR cells treated with water or LPS (500 ng/mL) were separated on SDS-PAGE and probed for p-TBK1, total TBK1, and MNRR1. Actin was probed as loading control. **(B)** Placental lysates from control (PBS) versus LPS (intraperitonially) injected mice were separated on an SDS-PAGE gel and probed for NOX2 and ATM kinase levels. Actin was used as a loading control. **(C)** Mouse liver lysates from TLR4^−/−^ control (PBS) versus LPS injected mice were separated on an SDS-PAGE gel and probed for NOX2 and ATM kinase levels. Tubulin was probed as a loading control. **(D)** Equal amounts of WT or TLR4^−/−^ mouse macrophage cells were treated with the NOX2 inhibitor with or without LPS (500 ng/mL) and lysates were separated on SDS-PAGE and probed for MNRR1. Actin was probed as loading control. **(E)** *Left*, Intact cellular oxygen consumption in HTR cells overexpressing EV or CHCHD4 and treated with control (water) or LPS for 24 h. Data are expressed relative to EV-control set to 100%. *Right*, Pooled lysates from OCR measurement were separated on an SDS-PAGE gel and probed for CHCHD4 levels. Actin was probed as a loading control.

